# Development of rat organoids to study intestinal adaptations after Roux-en-Y Gastric Bypass

**DOI:** 10.1101/2024.02.24.581868

**Authors:** Soukaïna Benhaddou, Lara Ribeiro-Parenti, Amélie Vaugrente, Alexandra Willemetz, Claire Gaudichon, Véronique Douard, Maude Le Gall, Pierre Larraufie

**Affiliations:** Université Paris-Saclay, AgroParisTech, INRAE, UMR PNCA, 91120, Palaiseau, France; Inserm UMRS 1149, Centre de Recherche sur l’Inflammation; Université Paris Cité, 75018, Paris, France; Service de Chirurgie Digestive Oesogastrique et Bariatrique, Hôpital Bichat—Claude-Bernard, Assistance Publique-Hôpitaux de Paris, Université Paris Cité, 75018 Paris, France; Université Paris-Saclay, INRAE, AgroParisTech, Micalis Institute, 78350, Jouy-en-Josas, France

## Abstract

Organoids from intestinal regions have proven to be useful tools to study intestinal epithelial responses to different conditions. Roux-en-Y gastric bypass (RYGB) has been associated with important intestinal adaptations, but the mechanisms underlying these changes are still poorly understood. Organoids could therefore be used to better decipher the intestinal adaptations associated with this surgery. Rat is a common model to assess RYGB responses *in vivo*, but surprisingly, very few studies managed to develop organoids from rat small intestine. The primary objective of this study was to establish a protocol for cultivating organoids derived from the small intestine of healthy rats. The second objective of this study focuses on the development of organoids from the small intestine of rats subjected to RYGB to evaluate whether phenotypic or gene expression differences emerge.

We successfully devised a functional protocol for developing organoids from fresh or frozen rat small intestine tissues. The obtained organoids exhibit significant variability, making interpretation challenging. Variability is observed in size, shape, and the number of organoids developed from the same sample, but also gene expression, depending on samples prepared on different days or from fresh or frozen tissues. This protocol was then applied to the small intestine of RYGB or sham-operated rats. However, we did not detect any major difference in size between intestinal organoids derived from Sham rats and those from RYGB rats. The expression of several genes (peptide transporters, amino acid transporters, genes specific to certain types of intestinal cells, etc.) was also assessed, and inter-experiment variability was higher than any effect due to the operation on the rat the intestinal tissue was originating.

In conclusion, this study has established a functional protocol to grow small intestine organoids in rats. Initial results suggest that in our experimental conditions, organoids obtained from rats subjected to RYGB do not differ from those obtained from Sham rats. However, increasing the sample size and improving reproducibility between experiments will be essential to confirm these findings.

## Introduction

Bariatric surgery is one of the most effective treatments for morbid obesity that is associated with long-term metabolic beneficial effects. Among them, Roux-en-Y-Gastric Bypass (RYGB) is the second most commonly performed bariatric surgery worldwide (29% of bariatric surgeries) after sleeve gastrectomy (63%). The bypass of most of the stomach, the duodenum and a part of the jejunum, connecting the upper pouch with the jejunum, leads to the creation of three intestinal limbs and reduces, the length of the small intestine, as well as food intake and nutrient absorption capacities. The *biliopancreatic limb* receives pancreatic secretions and bile (corresponding to the duodenum / proximal jejunum), the *alimentary limb* allows the passage of the food bolus (corresponding to the intermediate jejunum), and the contents of these limbs flow together into the *common limb* (corresponding to the distal jejunum) [1]. RYGB produces many beneficial effects, such as sustained weight loss and improvements in comorbidities [2]. Several mechanisms have been proposed and are debated to explain these effects. Among them, elevated postprandial gut hormones rapidly occur after the operation and could participate in beneficial metabolic effects before weight loss-induced effects [3–5]. The increase in gut hormones can result from the altered nutrient flow of nutrients [6, 7] and other intestinal adaptations after RYGB.

Among intestinal adaptations, Stearns and collaborators showed that villus height and crypt depth were increased in the alimentary and common limbs of RYGB-operated rats [8]. This observation has been reproduced several times and confirmed in humans [9–13]. One of the consequences of this hyperplasia is the increased total number of enteroendocrine cells, the cells secreting gut hormones, and it has been suggested that this mechanism could partially explain the increased circulating gut hormone levels after surgery [14]. Intestinal hyperplasia is seen as an adaptation of the intestinal epithelium, increasing nutrient absorption in response to the malabsorption induced by RYGB. However, the mechanisms involved in this phenomenon and its consequences on physiology are unknown.

Understanding RYGB-induced intestinal adaptation has been limited by the invasiveness of the procedures required to assess intestinal development, by the severity of procedures on animals and by the complexity of factors regulating intestinal epithelium. The development of intestinal organoids has been seen as an opportunity to study the response of intestinal epithelium to specific conditions while reducing cofounding effects. Organoids are grown from stem cells located in the crypts of the epithelium and successfully reproduce intestinal epithelial development, differentiation, and growth in response to different environmental cues [15, 16]. Interestingly, organoids also retain some memory of the tissue from which they originate, such as their location along the gastrointestinal tract [17], or adaptations induced by pathologies [18]. Moreover, organoids can be used to assess *in vitro* the mechanisms regulating intestinal epithelial proliferation, differentiation and functions by changing, in a controlled manner, their culture environment, higher throughput for testing different compounds, and assaying responses by microscopy or molecular biology approaches. One limitation of the studying human organoids is obtaining the appropriate controls: it is often difficult to obtain samples before and after the RYGB and to determine which pre-surgery sampled tissue is the appropriate control for each post-surgery intestinal limb. The rats have been commonly used as a model for RYGB, since they recapitulate well the intestinal adaptation to RYGB surgery in humans and tolerate this type of surgery better than mice. Rats could thus overcome the limitations of the appropriate controls since sham-operated animals can be used to generate control-paired samples. Therefore, using rat RYGB model together with rat organoid studies is a promising approach to study the intestinal adaptation to gastric bypass surgery. Yet, despite organoids being developed from many animal models, very few studies developed organoids from rat small intestine [19–21].

We therefore developed a protocol to grow small intestine organoids derived from rats in order to compare organoid development from crypts isolated from RYGB-operated rats or control animals as we hypothesized that RYGB-induced intestinal rearrangements might alter stem cell programming to induce hyperplasia that can be reproduced in organoids. In this study, we report crypt isolation and organoid growth from different rat jejunal regions, including from frozen sample tissues and present some first results comparing the growth and expression of intestinal markers of organoids derived from RYGB-operated rats.

## Materials and Methods

### Animals

For organoid production development, intestinal samples from proximal and jejunal samples (∼2cm^2^) were obtained from euthanized male wistar rats housed at the IERP animal facility (agreement number C78-120). After euthanasia, intestines were rapidly sampled, intestinal content removed, and samples placed either in Leibovitz’s L-15 medium (Gibco 11415-064) for immediate use or slowly frozen in fetal bovine serum (FBS) with 10% DMSO and 10µM Y27632 (Biotechne 1254/10) in an isopropanol recipient and stored in liquid nitrogen.

Samples obtained to study the effect of bariatric surgery on organoid phenotype were obtained from rats included in a larger study about intestinal adaptation after bariatric surgery. This study complied with European Union Directive 2010/63/EU for animal experiments. It was validated by the Institutional Animal Care and Use Committee (Comité d’Ethique Paris Nord, n°121, experiment and animal facility approval number: 34483).

Female Wistar rats aged 7 weeks were housed under standard environmental conditions for 4 months at a maintained temperature of 21°C-22°C, a 12/12h light/dark cycle, and with water and food HFD (High-Fat Diet) available ad libitum.

### Surgical Procedures

The animals were randomly divided into two groups: RYGB (n= 5) and sham-operated (Sham, n= 5). They were anesthetized by the gaseous inhalation of isoflurane (Vertflurane, Virbac, France). The procedures were performed as previously described [12].

Rapidly, the surgical procedure started with a laparotomy, isolating the stomach outside the abdominal cavity and removing the non-glandular part of the stomach (forestomach) by applying a staple line. A gastric pouch was created using another staple line parallel to the first. The remaining gastric pouch corresponded to 20% of the initial stomach size. The jejunum was transected 20cm after the pylorus, the alimentary limb was anastomosed to the gastric pouch, and the biliopancreatic limb was anastomosed 15cm distally to the gastrojejunal anastomosis. For Sham-operated rats, an unarmed staple gun was used to pinch the stomach to mimic surgery.

Following all procedures, the laparotomy was closed using 4-0 vicryl and 3–0 vicryl (Ethicon) sutures to sew up the abdominal wall and the skin, respectively. An analgesic (Xylocaine, Astra, 10 mg/kg) was infiltrated along all the sutures to reduce pain, and an intramuscular injection of antibiotic (Penicillin, 20,000 U/kg, PanPharma, Boulogne Billancourt, France)) was administered. After 3 days of a liquid meal, the rats were switched back to the HFD.

Rats were euthanized 3 weeks after surgery. For each experiment, samples from one sham rat and one RYGB rat were processed in parallel to take into account day-to-day experimental variability. Two samples of the small intestine were collected: a sample of proximal jejunum and medium jejunum in Sham rats, and a sample of biliopancreatic limb and alimentary limb in RYGB rats. 6 rat samples (ExpA, B and C) were placed directly in Leibovitz’s L-15 medium (Gibco 11415-064) to develop organoids immediately. The samples of other rats were placed in a freezing medium composed of 90% FBS, 10% DMSO and 10µM Y27632 to be processed all at the same time (ExpD).

### Organoid growth media

For the culture of undifferentiated organoid cultures, we used a conditioned medium made from L-WRN cells (ATCC CRL3276), containing Wnt3a, Noggin, and R-Spondin as follow: After two weeks of cell-line selection using G418 and hygromycin, L-WRN cells were grown to confluency. Conditioned media was produced by adding fresh advanced DMEM/F12 media (Gibco 12634-010) with 2mM Glutamax and 10% FBS, collecting it, and replacing it four times every 24 hours. Collected conditioned media were stored at 4°C before being pooled, filtrated at 0.22µm, aliquoted, and stored at -80°C.

Growth rat organoid media was then produced by adding 50% (vol/vol) L-WRN conditioned media, 40% advanced DMEM-F12, 10% FBS with 50ng/mL EGF (Gibco PMG8041), 1x B27 (Gibco A3582801), 1x N2 (Gibco A1370701), 1mM N-acetyl-cystein (Merck A9165), 100ng/mL IGF-1 (Preprotech 100-11), 100ng/mL FGF-2 (Preprotech 100-18B), 500nM A8301 (Preprotech 9094360), 10nM Gastrin (Merck G9145) and penicillin (50IU/mL) and streptomycin (50µg/mL).

Differentiation media for rat organoids was composed of 10% Noggin-conditioned media, produced from 7-days supernatant of a confluent culture of HEK293 – mouse Noggin -FC in advanced DMEM-F12, 90% advanced DMEM-F12, 50ng/mL EGF, 1x B27, 1x N2, 1mM N-acetyl-cysteine, 100ng/mL IGF-1, 100ng/mL FGF-2, 500nM A8301, 10mM Gastrin and 10mM DAPT (Biotechne 2634).

### Stem cell isolation and culture

Intestinal crypts were isolated similarly from tissue from the day of euthanasia (kept at 4°C) or from tissue rapidly thawed after liquid nitrogen storage. Briefly, 2 cm^2^ of each sample was collected, flushed, and washed several times with ice-cold calcium/magnesium-free PBS. Then, the mucus and villi were removed by gently scraping the opened tissue with flat tweezers. The tissue was then cut into 5mm pieces, placed in 5mL of ice-cold PBS, and shaken vigorously. The supernatant was removed, and 5mL of Gentle Cell Dissociation Reagent (StemCells 100-0485) was added to the tube. The tube was incubated on ice for 20 min with gentle agitation every 10 minutes. The Cell Dissociation reagent was replaced with ice-cold PBS, and the tissue was vigorously triturated with a 5mL pipette several times. The tissue was allowed to settle, the supernatant was collected, and 5mL of fresh PBS was added. Those steps were repeated until fragments from crypts without villi debris could be observed under the microscope (Fig1A). The cell solution was then supplemented with 1mL of advanced DMEM-F12 with 10% FBS and penicillin/streptomycin and centrifuged at 300g for 4 minutes at 4°C. The pellet of intestinal crypts was resuspended in 100 µL of Matrigel (Corning 356235), and domes of 10µL placed in 12-well plates (3-4 domes per well). Matrigel was set for 30 minutes at 37C before adding 500µL organoid growth media with 10µM Y27632. Media was replaced after 24h with organoid growth media without Y27632 and changed every other day. Round organoid structures begin forming within 24-48 hours.

After 6-8 days of culture, organoids could be passed when dead cells accumulated in the organoid lumen. The medium was removed, and cold DMEM F12 medium was added to dissolve the matrigel domes. The cell suspensions were pulled together and centrifugated at 150g for 4 minutes at 4°C to eliminate the cell debris. The supernatant was removed, and organoids were incubated in 300µL TrypLE (Gibco 12505010) at 37°C for 2 min. Then, 700µL of DMEM F12 medium containing 10%FBS was added and organoids spilt by trituration. After centrifugation at 250g for 4 minutes at 4°C, the supernatant was removed, the pellet was resuspended in Matrigel, and new domes were seeded in 24well plates. After 30 minutes for the domes to set, organoid growth media was added with 10µM Y27632 for the first 48 hours, and the media changed every other day.

For differentiation experiments, organoids were grown for 4 days in organoid growth media, and then the media was replaced with organoid differentiation media for 48 hours.

Organoids were regularly pictured using a microscope (Olympus BX43) to validate organoid growth and measure their size using NDP.view2 software. Statistical analysis was performed using individual organoid size from all samples, but experiment variability was taken into account using a two-way factor ANOVA using the experiment day as the second factor. Hierarchical clustering was performed using the pheatmap R package.

### RNA Extraction, Reverse Transcription, and qPCR

After 8 days of culture, organoid media was removed, and cold advanced DMEM-F12 was added to wells to harvest organoids. Organoids were centrifuged at 300g and 4°C to remove the wash media, lysed in 300µL of RLT buffer (Qiagen) and stored at -80°C until extraction of all samples. During crypt isolation, the fraction of villi and crypts were also isolated, rinsed in PBS, and lysed in 300µL of RLT buffer for comparison with organoids. RNA was extracted using a Qiagen RNeasy micro kit using the supplier’s instruction and eluted in 14µL of RNase-free water. RNA concentration was measured using a nanodrop, and 200ng of RNA was reverse-transcripted using a High-Capacity cDNA Reverse Transcription Kit (Applied Biosystems, 4368814) in 20µL.

Gene expression was measured by qPCR using a StepOne Plus Real-Time PCR system (ThermoFisher Scientific) in 12µL reactions with 6µL of PowerUp SYBR Green Master Mix (Applied Biosystems A25741), 0.2µL of cDNA sample and 200nM of primers (Table 1). Gene expression was quantified based on Ct evaluation using the StepOne Software v2.3 (ThermoFisher Scientific) and expressed relative to the mean expression of *18S* and *Tbp* using the 2^-ΔCt^ method. Differential expression between samples was assessed using a two-way ANOVA to take into account the day of experiment in addition to the group of origin of the tissue.

**Table 1.**
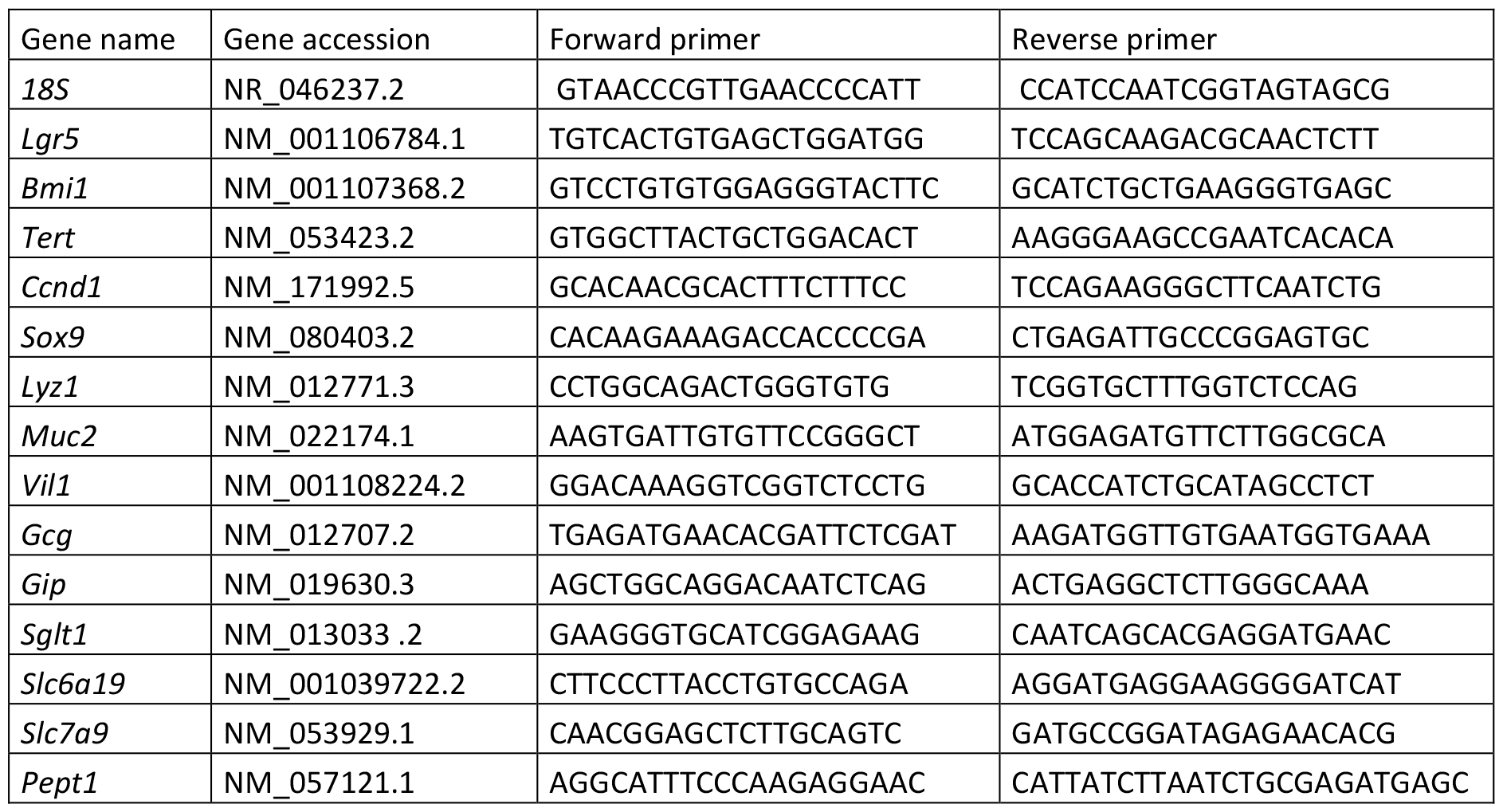
Genes and primers.

## Results

### Development of rat jejunal organoids

As no method to grow organoids from rat jejunum was published when we developed this protocol, we first tested if methods used for generating human or mouse organoids would work. We tested standard protocol with crypt isolation by tissue trituration or scrapping after incubation with different EDTA concentrations (2 to 20mM) followed by growth in mouse organoid culture composed of conditioned media from L-WRN cells mixed with one volume of advanced DMEM-F12 and 10 % FBS or human growing condition, either using human Intesticult growth media (StemCells) or the mouse media complemented with B27, N2, 1mM N-acetyl-cystein, 100ng/mL IGF-1, 100ng/mL FGF-2, 500nM A8301, 10nM Gastrin and 10mM nicotinamide. No organoids grew sustainably in these conditions, with only very few crypts forming small organoid-like structures that did not develop. We, therefore, adapted our method to isolate crypts from proximal and intermediate jejunum using StemCells Gentle Cell dissociation reagent instead of EDTA and removed nicotinamide from the human-conditioned media just presented. This method resulted in a relatively similar ratio of debris (Figure 1A). However, after 3 days, several organoid spheres could be observed in more than 75% of tested samples (Figure 1B). Then, they could grow into bigger organoids and some, but not all developed the budding characteristics of organoids (Figure 1C). Of note, organoid growth was very variable within one sample, but also from one preparation to the other. After 7-10 days, organoids blackened and died. Passage of organoids after 6-8 days enabled the proliferation of organoids. We successfully passaged organoids for up to 3 passages (we did not test for longer) and differentiated them using a media without R-spondin and Wnt3a but with DAPT. We confirmed organoid differentiation by qPCR by looking at the expression of different cell markers (Figure 1D) and comparing it to isolated crypts. As expected, expression of proliferating cell markers (*Lgr5, Bmi1, Tert*) and early differentiating cells (*Sox9*) was higher in undifferentiated organoids, and differentiated cell markers such as *Muc2, Gcg, Gip, Sglt1* or *Slc6a16* were higher in differentiated organoids than undifferentiated ones. Whereas, we did not manage to freeze crypts and grow them afterwards, organoids could be produced from tissue that has been frozen before isolation of crypts. This enables one to use material from different experiments at the same time thus likely reducing variability even if organoids from frozen samples grew smaller during the first passage. We also could successfully grow organoids from female and male rats and we did not notice any difference between both sexes.

**Figure 1.**
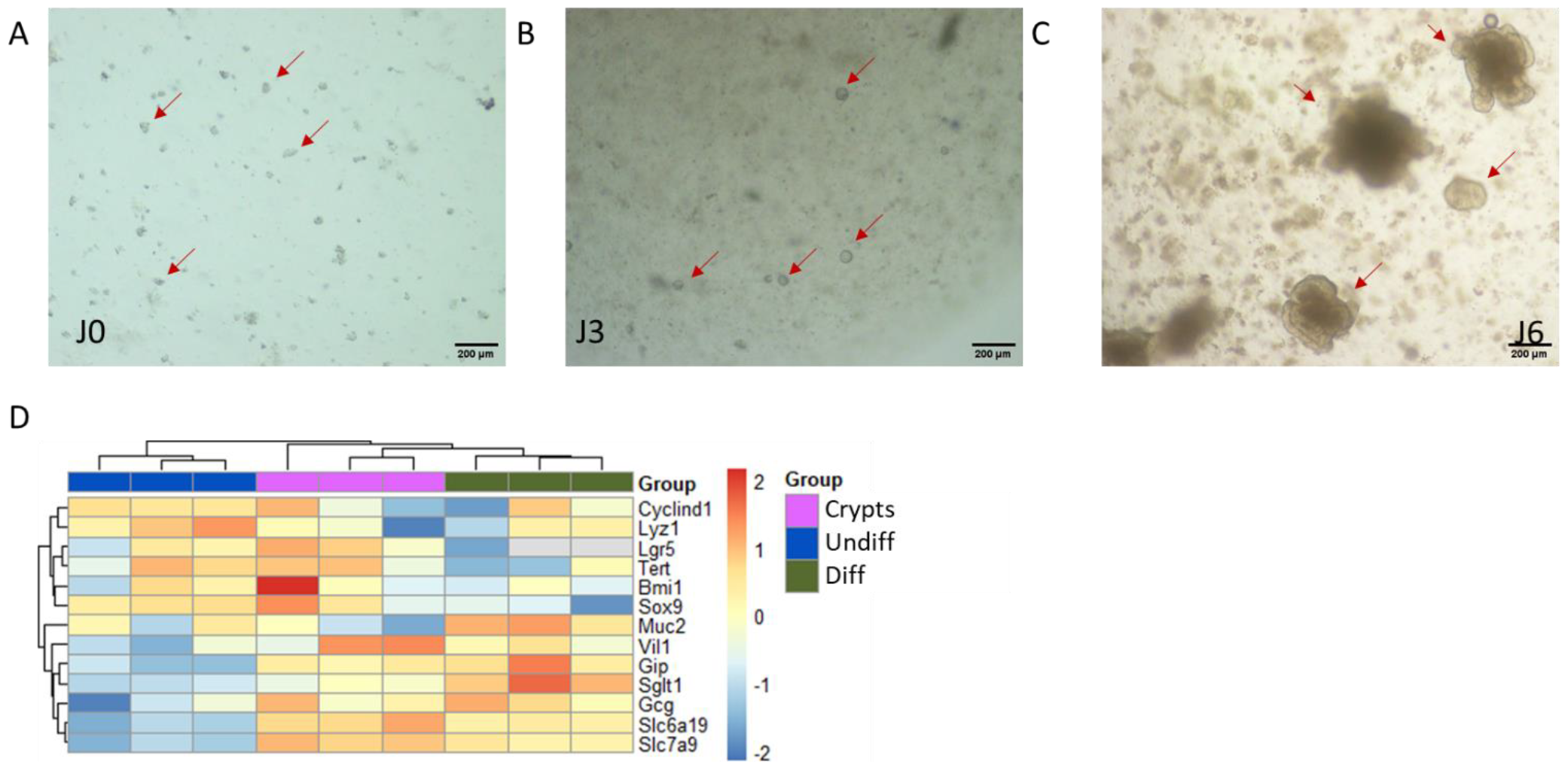
Establishment of rat small intestine organoid from a fresh biopsy of normal tissue. Microscopy images of crypt isolation (A), development of organoids after 3 days of culture (B), and 6 days of culture (C). Arrows indicate crypt and organoid structures. (D) heatmap representing relative gene expression normalized to the mean of the two housekeeping genes, 18S and TBP, and overall expression across all samples. Samples from isolated crypts are highlighted in purple, undifferentiated organoids (Undiff) in blue and differentiated organoids (Diff) in green.

### Growth of organoids issued from RYGB or sham-operated rats

Using our developed protocol, we isolated crypts from sham- and RYGB-operated rats on fresh and frozen tissues, always pairing samples between an RYGB and a sham and grew organoids. The biliopancreatic limb was compared to the proximal jejunum and the alimentary limb to the intermediate jejunum of sham rats. Six days after crypt isolation, the organoids were counted, and their size was analyzed by measuring the average diameters in both directions. Microscopy images show that the number of organoids obtained varies significantly from one set of experiment to the other. Moreover, for each set of experiments, the size and shape of organoids were highly variable within a condition, with a coefficient of variation within each condition ranging from 40 to 58%. Indeed, some organoids looked more like spheroids, while others developed many buds, but we did not investigate if these differences represented a differentiation process (Figure 2A-D). Notably, the organoids developed from fresh tissues (Figure 2A-D) seemed bigger than those derived from frozen tissue (Figure 2E-F). When comparing experiments, we did not observe any major and significant difference in the size of organoids from Sham and RYGB rats in both regions (Figure 2G-H). However, the high variability between experiments suggests that more experiments would be required to detect a slight change in organoid size. In order to take into account the high variability between experiments day, we used a two way ANOVA analysis (Table 2). In both intestinal regions, the effect of the day of treatment was significant, highlighting the difficulty to grow similar organoids on different experiments. However, as we counted many organoids from each experiment and condition, the general effect size was relatively low for the effect of the surgery on organoid development, indicating that in these conditions, we could not detect an effect of the surgery on organoid growth.

**Table 2:**
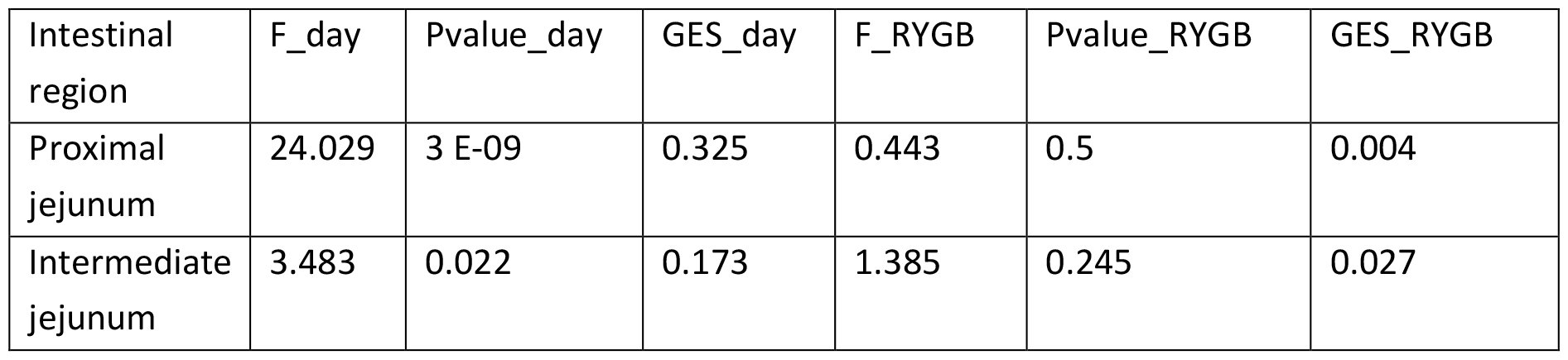
summary of ANOVA results from testing the effect of experimental day and surgery on organoid size, each single organoid being considered as an independent sample.

**Figure 2.**
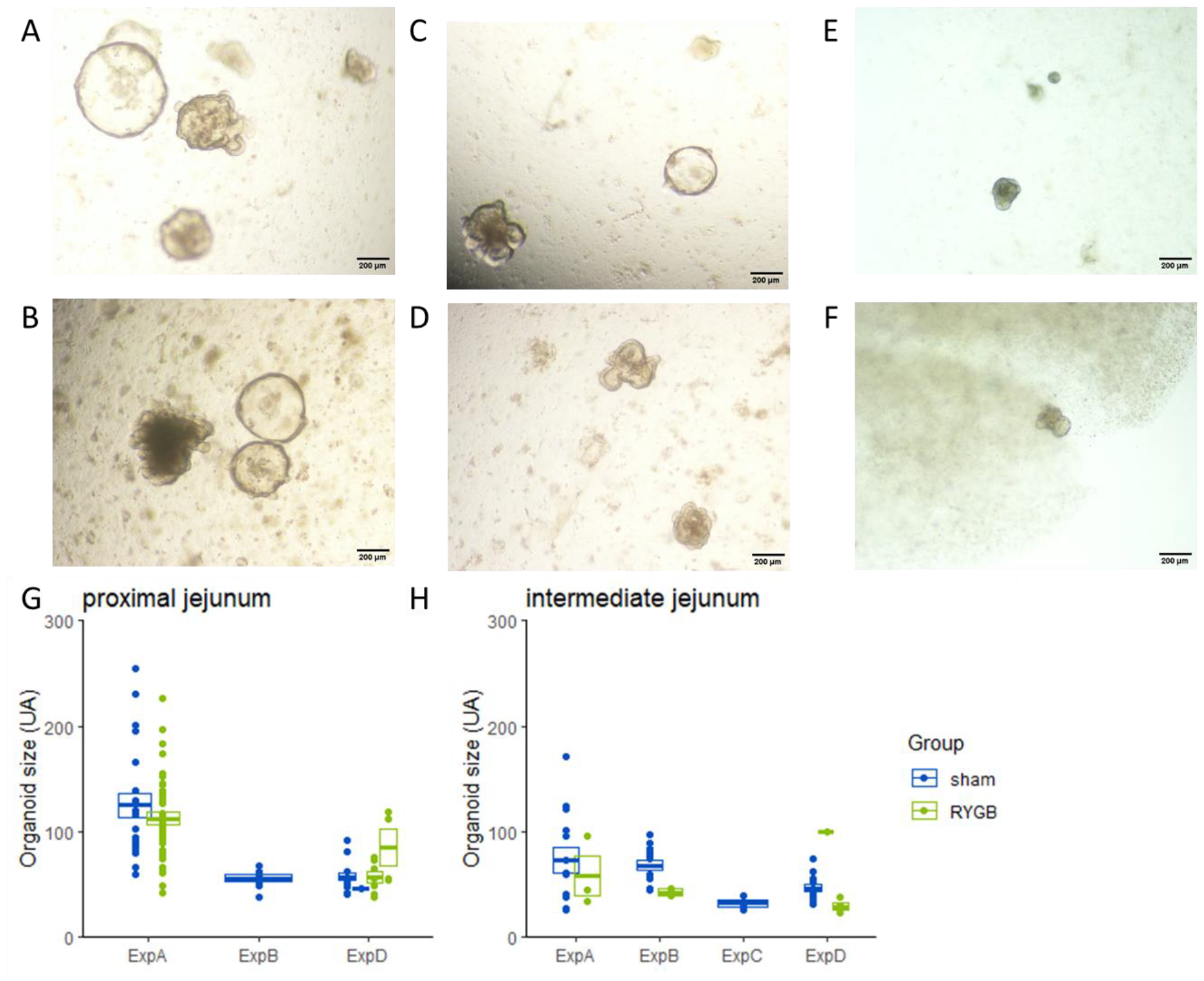
Number and size of organoids generated from Sham rats and RYGB rats Representative images of organoids developed from Sham rats (A, C, E) or RYGB (B, D, F) rats in the proximal jejunum (corresponding to the biliopancreatic limb in RYGB rats) (A and B) and the intermediate jejunum (corresponding to the alimentary limb) (C-F) in fresh (A-D) or frozen tissues (E, F)). Size of organoids measured in the proximal (G) and intermediate jejunum (H) in Sham rats (blue) and RYGB rats (green). Different sets of experiments are indicated on the x-axis for intra-experiment comparison, with individual organoid size shown for each experiment and condition, with mean ± se depicted.

### Expression of proliferation and differentiation factors

We assessed potential modifications of organoid differentiation or change in nutrient absorption activity by measuring expression of genes involved in these functions. RNA was extracted from organoids generated from proximal and intermediate jejunum of Sham and RYGB rats after 6 days of culture. RT-qPCR were performed on key genes involved in cell proliferation, epithelial cell differentiation and genes involved in nutrient transport and markers of intestinal epithelial cell populations (Figure 3). We observed a very high variability in gene expression between samples from different set of experiments within a same condition, as the mean coefficient of variation for all genes tested was 105%. However, organoids derived from Sham and RYGB operated rats in both segments of the small intestine had similar average gene expression but the low number of experiments greatly and high variability limited the scope of the conclusion we could draw from these experiments.

**Figure 3.**
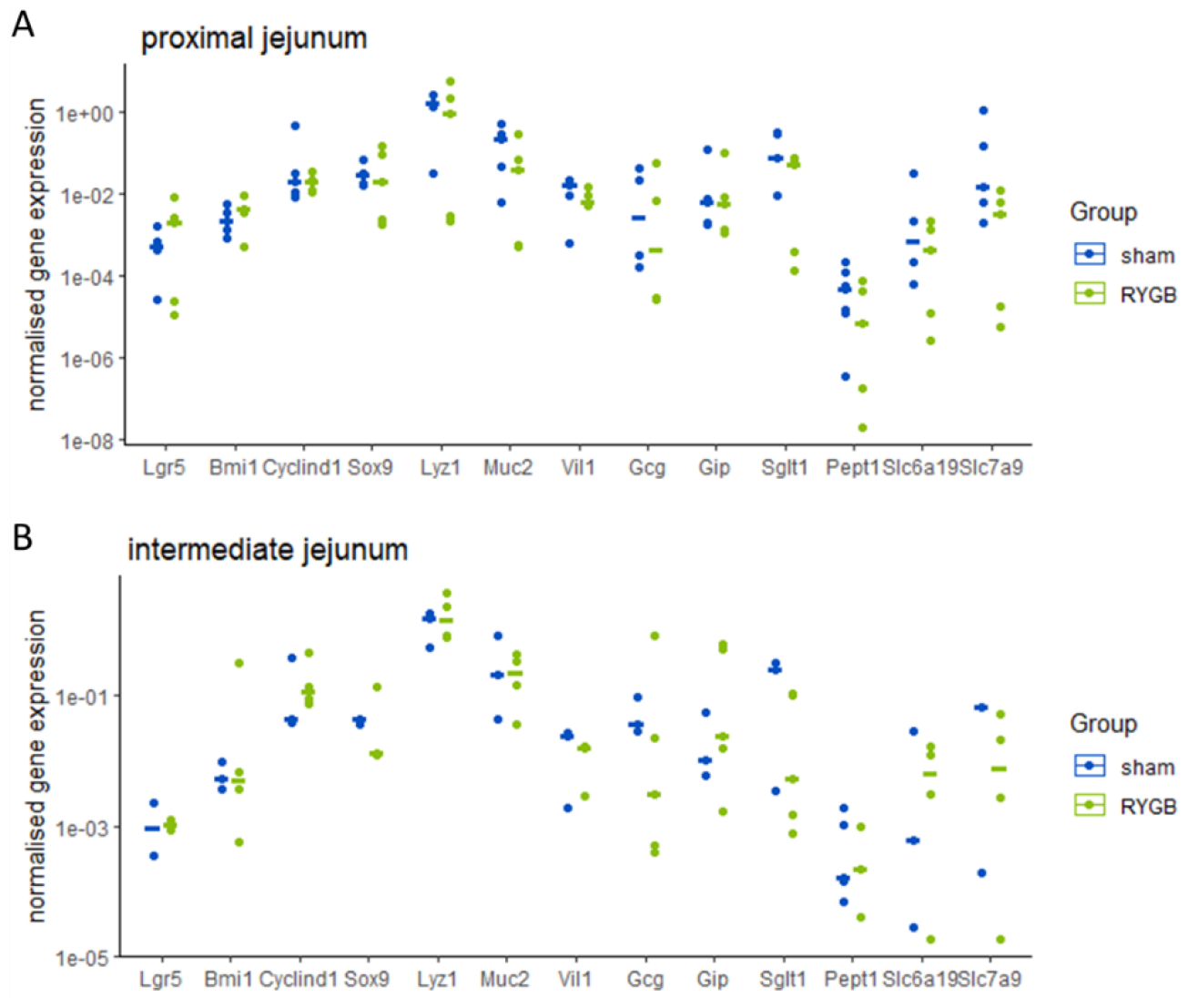
Relative mRNA expression of genes Normalized gene expression in organoids derived from proximal (A) and intermediate (B) jejunum of Sham rats (blue) and RYGB rats (green).

## Discussion

To our knowledge, this study represents the first attempt at growing organoids derived from jejunum tissues from rats operated on bariatric surgery. Even if the method is not as efficient as growing organoids from mouse or human samples, after testing different conditions, we successfully obtained organoids from most of the samples we processed, with a 75% success rate. The method we present can thus be helpful to test intestinal adaptation *in vitro* in rats, such as assessing the effect of RYGB surgery on intestinal adaptation. Similar to organoids from other species, rat organoids showed high size and shape variability, both within the same sample and between experiments performed on different days. The possibility to freeze tissue seems to be an advantage to overcome this variability issue. However, we observed a lower success rate and a reduced organoid size, indicating a potential effect of freezing on cell survival and possibly stem cell programming. In our hands, crypt isolation was more sensitive in rats than in other species and required an adapted protocol to increase the viability rate. The media we used for growing rat organoids was closer to human ones than the mouse ones as it required IGF-1 and FGF-2. Even if we did not test the importance of all compounds in organoid growth, FGF2 seemed to be a key factor for the growth of rat organoids, whereas IGF1 seemed less important even if we kept it in our experiments.

This model of organoids derived from rats was then used to assess if the intestinal hyperplasia induced by RYGB surgery is passed by stem cells and can be reproduced *in vitro*. This hyperplasia may be explained by the drastic change of environment encountered by intestinal epithelium due to the rearrangement of the surgery, the change in microbiota composition, inflammation in the tissue or the change in nutrient flow after surgery. However, the effect on cell proliferation could either come from a direct sensing of the environment by proliferative cells or an imprinting of stem cells changing their proliferation and differentiation profile. To test the latter hypothesis, we isolated stem cells form RYGB operated rats and grew organoids from these cells as we would expect that organoids derived from these stem cells could grow larger. We observed a high variability between samples within each group that limited our capacity to detect smaller size effects, however these experiments indicated that there was not a large size effect in organoid development from crypts isolated from RYGB-operated rats compared to those from sham-operated rats.

Moreover, the organoids expressed similar levels of gene markers for different intestinal cell populations within the high variability within groups, indicating that effect of organoid growth variability due to experimental conditions was higher than the potential effect of the origin of the sample. However, due to the high variability of organoid growth, we could not exclude a smaller effect kept by stem cells, and more replicates of these experiments would be needed to assess these effects more precisely. Moreover, we did not assess the importance of the change of environment induced by RYGB surgery. Organoids were grown in a nutrient-rich media that does not represent the environment the intestinal cells encounter. This media could reduce the adaptation of stem cells to their RYGB environment *in vivo* as all organoids had access to similar and high concentrations of nutrients and factors, therefore possibly modulating cell development and masking the effect driven by stem cells. Modulating nutrient availability in the media or other factors affected by the surgery could therefore be key to reproduce *in vitro* intestinal hyperplasia. Indeed, it can be hypothesized that jejunal epithelial cells from RYGB operated rats have access to more nutrients due to lower absorption. One hypothesis to explain intestinal hyperplasia is based on the action of GLP-2, which is released in high quantities after food intake and has an intestinotrophic action. However, GLP2 action does not seem direct on intestinal cells as intestinal cells do not express GLP2 receptors. The role of fibroblasts secreting IGF-1 in response to GLP-2 could be key [22, 23]. However, in our hands, the IGF-1 concentration did not seem to influence rat organoid growth, but further studies would be required to precisely assess its role by controlling precisely organoid growth and by reducing the quantity of serum used in this organoid growth as calf serum contains several growth factors that could interfere with the analysis.

## Notes

### Competing Interest Statement

The authors have declared no competing interest.

